# Tissue distribution and tumor concentrations of hydroxychloroquine and quinacrine analogs in mice

**DOI:** 10.1101/496018

**Authors:** Abigail R. Solitro, Jeffrey P. MacKeigan

**Author notes:** **CORRESPONDING AUTHOR** (JPM).

## Abstract

Hydroxychloroquine (HCQ) is a 4-aminoquinoline molecule used for the treatment of malaria, and more recently to treat rheumatoid arthritis, systemic lupus erythematosus, and cancer. In cancer, HCQ is being used in multiple cancer clinical trials as an inhibitor of autophagy, a cytosolic degradation process employing the lysosome. Importantly, more potent lysosomotropic agents are being developed as autophagy inhibitors. Additional studies revealed that acridine-based compounds such as quinacrine (QN) increased potency over the 4-aminoquinoline HCQ. In line with these initial discoveries, we performed chemical synthesis of acridine-based compounds and screened for potent autophagy inhibition. The novel compound VATG-027 increased potency and cytotoxicity over HCQ in osteosarcoma and melanoma cell lines, supporting further investigation in vivo. Here, we developed a liquid chromatography tandem mass spectrometry (LC-MS/MS) method to investigate HCQ, QN, and VATG-027 compound concentrations across various tissue types in mice. This method detected compound concentrations in whole blood, lung, liver, kidney, and subcutaneous tumor tissues. Concentrations of HCQ, QN, and VATG-027 varied within and between tissue types, suggesting unique tissue distribution profiles for 4-aminoquinoline and acridine compounds.

## INTRODUCTION

Hydroxychloroquine (HCQ) is a 4-aminoquinoline molecule initially approved by the Food and Drug Administration (FDA) for the treatment of malaria (1, 2). The lysosomotropic characteristics of HCQ render erythrocyte lysosomes inactive, effectively starving *Plasmodium* parasites and eliminating these disease-causing organisms (3–5). HCQ is also used to treat various autoimmune diseases, including rheumatoid arthritis and systemic lupus erythematosus (6–9). Further, HCQ is being used in cancer clinical trials as an inhibitor of macroautophagy, a cytosolic degradation process employing the lysosome (10–12).

Macroautophagy (hereafter, autophagy) has been implicated as a chemoresistance mechanism, as it sustains cancer cell viability in response to stresses such as nutrient starvation, hypoxia, and cytotoxic therapies (13–17). In addition, mutations in prominent proto-oncogenes, including *KRAS* and *BRAF,* elevate autophagy in cancer cells inducing a state of autophagy dependence (18–23). These studies accelerated the clinical investigation of HCQ as an adjuvant to combination therapy strategies. To date, 13 clinical trials have evaluated HCQ as an anti-cancer agent (24–36). Although HCQ has achieved partial responses and stable disease in some cases, the compound lacks potency, producing variable degrees of autophagy inhibition across patients receiving the maximum tolerated dose of 1200 mg per day (30, 32). As a result, more potent lysosomotropic agents are being developed (37). Early studies demonstrated that dimeric compounds based on the 4-aminoquinoline structure were more potent than the monomeric HCQ (38). Additional studies revealed that acridine-based compounds such as quinacrine (QN) increased potency over the 4-aminoquinoline HCQ (39, 40). In line with these initial discoveries, we performed chemical synthesis of acridine-based compounds and screened for potent autophagy inhibition (39, 40). The novel compound VATG-027 increased potency and cytotoxicity over HCQ in osteosarcoma (U2OS) and melanoma (A375, UACC-91, UACC-257, UACC-502, UACC-903, UACC-1308, UACC-1940, UACC-2534, and UACC-3291) cell lines, supporting further investigation *in vivo* (39).

We developed a liquid chromatography tandem mass spectrometry (LC-MS/MS) method to investigate HCQ, QN, and VATG-027 compound concentrations across various tissue types in mice. This method detected compound concentrations in whole blood, lung, liver, kidney, and subcutaneous tumor tissues with a lower limit of quantification (LLOQ) of 2.5 ng/mL. Concentrations of HCQ, QN, and VATG-027 varied within and between tissue types, suggesting unique tissue distribution profiles for 4-aminoquinoline and acridine compounds. Additionally, our observations regarding tissue distribution will inform *de novo* drug development and provide insights into potential adverse side effects.

## RESULTS

The lysosomotropic compounds HCQ and QN are used to treat malaria and autoimmune diseases, and clinical trials are currently investigating HCQ as an anti-cancer adjuvant (1, 6, 10). The known lysosomotropism of these compounds led us to synthesize VATG-027, an acridine-based compound with increased potency and cytotoxicity over HCQ in osteosarcoma, melanoma, and NSCLC cell lines (Supplementary Figure 1) (39–41). In Figure 1, we compare the chemical structures of HCQ, QN, and VATG-027. The chemical structures produced unique physiochemical properties among the three compounds such as solubility and physical state, which are summarized in Supplementary Table 1.

**Figure 1.**
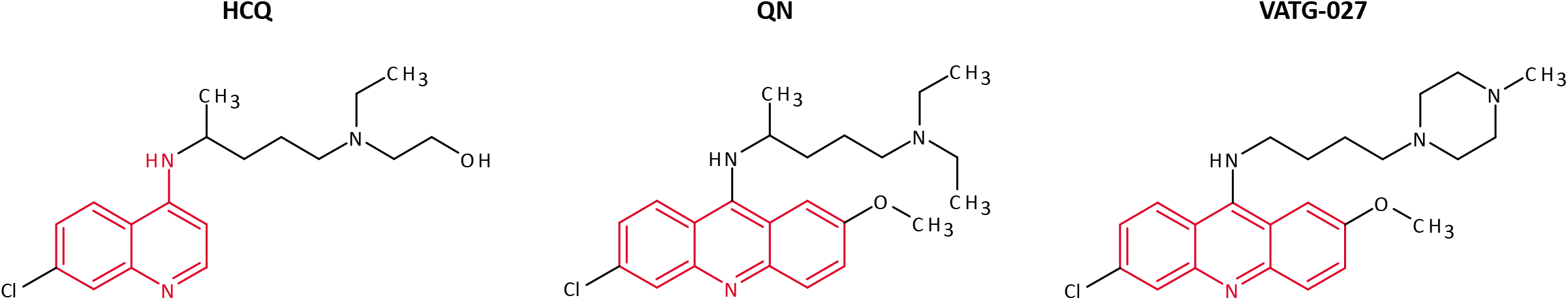
Chemical structures of lysosomotropic compounds. Chemical structures of hydroxychloroquine (HCQ), quinacrine (QN), and VATG-027 are shown with heterocyclic nuclei highlighted in red (4-aminoquinoline for HCQ and acridine for QN and VATG-027).

**Supplementary Figure 1. Acridine compounds increase potency and cytotoxicity over HCQ.** (A) A549 NSCLC cells were treated with HCQ, QN, or VATG-027 for 24 h. Potency regarding autophagy inhibition was determined by the effective concentration, or the concentration producing LC3B-II protein accumulation over vehicle treatment. (B) A549 cells were treated with HCQ, QN, or VATG-027 for 24 h. Viability measurements were compared to vehicle controls to produce relative viability measurements. Orange data points indicate the half-maximal effective doses.

To determine the *in vivo* characteristics of VATG-027 as compared to HCQ and QN, we developed an original LC-MS/MS method. Using reversed-phase high performance liquid chromatography (HPLC), we purified compound structures on a Phenomenex Luna C18 (2) column using gradients of two mobile phases, including mobile phase A (100% v/v water, 0.05% v/v formic acid, and 0.05% trifluoroacetic acid) and mobile phase B (50% v/v acetonitrile and 50% v/v methanol) (Table 1). Immediately following, we performed tandem mass spectrometry using a 12-point standard curve. To account for instrumental variability, we used a quinacrine mustard internal standard in each standard and experimental sample. Quadratic models were then fit to the standard curves and used to determine compound concentrations from peak area ratios, defined as the peak area of the compound relative to the peak area of the internal standard.

**Table 1.** Mobile phases in HPLC.

| Time (min) | % Mobile phase A | % Mobile phase B |
| --- | --- | --- |
| 0 | 85 | 15 |
| 0.1 | 85 | 15 |
| 1.0 | 20 | 80 |
| 3.5 | 20 | 80 |
| 3.6 | 85 | 15 |
| 5.0 | 85 | 15 |

Mobile phase A (100% v/v water, 0.05% v/v formic acid, and 0.05% trifluoroacetic acid) and mobile phase B (50% v/v acetonitrile and 50% v/v methanol).

### Whole blood levels over time after a single dose

Using this LC-MS/MS method, we investigated the whole blood concentrations of HCQ, QN, and VATG-027 over time after a single dose. Athymic nude mice were treated with a single dose of either 30 or 60 mg/kg HCQ, QN, or VATG-027 by oral gavage. At 3, 6, 24, 48, and 168 h after gavage, we retrieved retro-orbital whole blood samples and performed LC-MS/MS. Figure 2 shows the experimental samples plotted against the standard curves and quadratic models (A for HCQ, B for QN, and C for VATG-027). The quadratic models are summarized in Supplementary Table 2.

**Figure 2.**
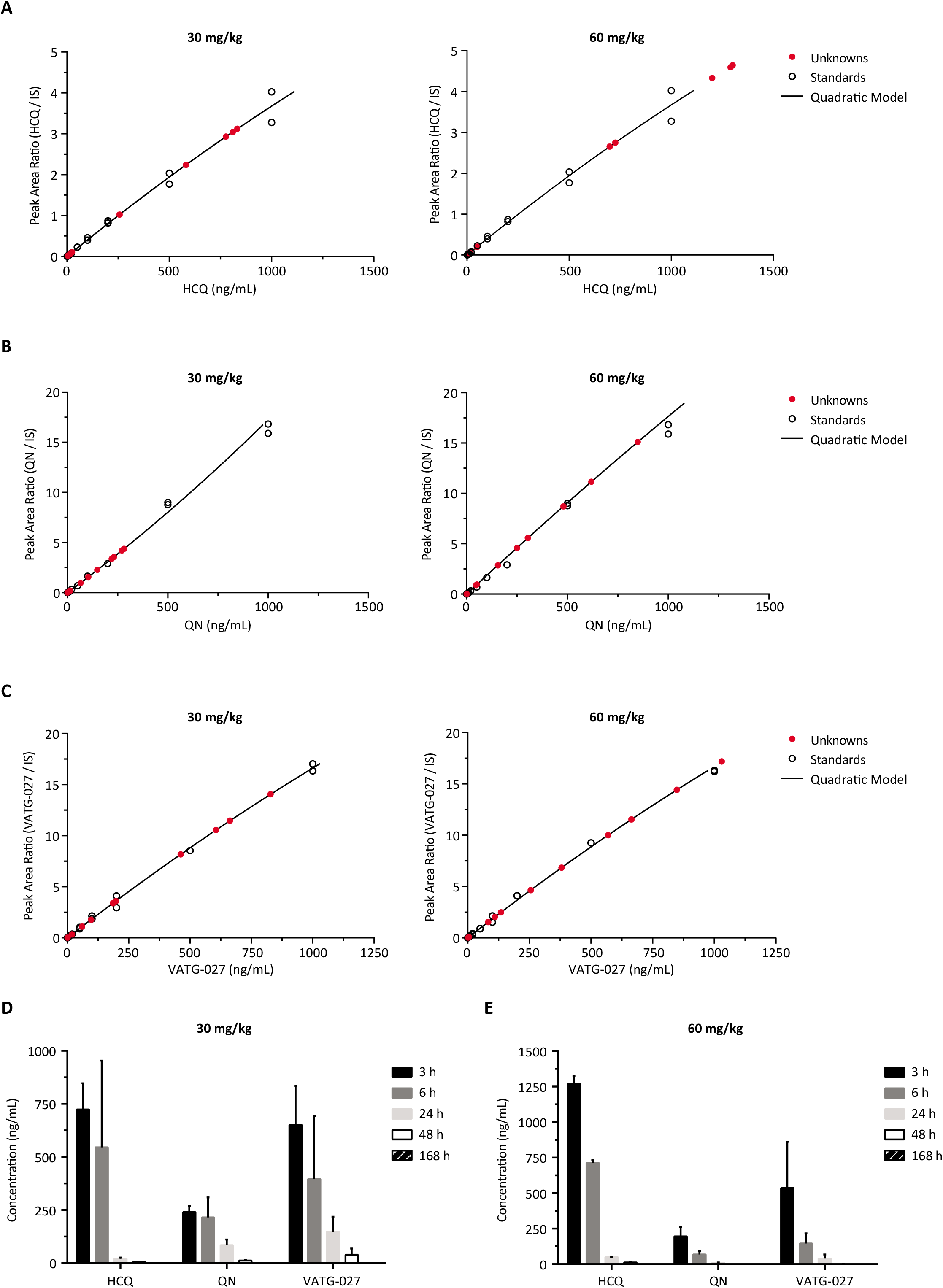
Whole blood levels over time after a single dose. (A) HCQ, (B) QN, and (C) VATG-027 experimental samples (red circles) are plotted against standards (open circles) and quadratic models fitted to the standards. (D) Athymic nude mice received a single dose of 30 mg/kg or (E) 60 mg/kg HCQ, QN, or VATG-027 by oral gavage and whole blood samples collected after 3, 6, 24, 48, or 168 h.

After a single 30 mg/kg dose, HCQ concentrations in whole blood were highest between 3 and 6 h, with mean concentrations at 720 and 550 ng/mL, respectively, for those timepoints (Figure 2D). Whole blood concentrations of HCQ significantly decreased from 6 to 24 h, with a mean 24 h concentration of 20 ng/mL. By 48 h post-gavage, HCQ concentrations were just above the LLOQ of this LC-MS/MS method. QN concentrations differed from HCQ: mean concentrations of 240, 220, and 80 ng/mL were measured at 3, 6, and 24 h post-gavage, respectively (Figure 2D). QN concentrations were slightly higher than HCQ concentrations after 48 h, but were also near the LLOQ. VATG-027 mean concentrations were 650, 400, 150, and 40 ng/mL for the 3, 6, 24, and 48 h timepoints (Figure 2D). Only after 168 h (or 7 d) were VATG-027 concentrations too low to detect in whole blood.

After a single 60 mg/kg dose, mean HCQ concentrations in whole blood were 1300, 710, 50, and 10 ng/mL at 3, 6, 24, and 48 h post-gavage, respectively (Figure 2E). Mean QN concentrations were lower at 200, 70, and 10 ng/mL at the 3, 6, and 24 h timepoints (Figure 2E). VATG-027 mean concentrations were also lower than HCQ, and were measured at 540, 150, and 40 ng/mL at 3, 6, and 24 h post-gavage (Figure 2E). Compound concentrations were undetectable after 168 h. These data indicate that whole blood concentrations are highest for the 4-aminoquinoline HCQ when compared to the acridine-based compounds QN and VATG-027. In addition, doubling the dose of each compound did not increase the whole blood concentrations by the same magnitude, suggesting efficient clearance from the blood.

### Whole blood and tumor concentrations after daily dosing

Lysosomotropic agents, such as HCQ, are being tested in combination treatment strategies for cancer (10). These agents may be particularly effective in the genetic context of KRAS-mutant NSCLC (18, 19). To investigate this, we established a xenograft mouse model using A549 NSCLC cells that contain a p.G12S mutation and treated with HCQ, QN, or VATG-027. A549 cells were subcutaneously implanted into the right flanks of athymic nude mice. After tumors grew to 150 mm^3^, mice were randomly assigned treatment groups receiving 30 mg/kg daily doses of HCQ, QN, or VATG-027 for 7, 14, or 28 d. Because plasma concentrations peak between 1 to 3.5 h after ingestion (42), we sampled whole blood and tumor tissue 3 h after the last dose for each treatment group. We then performed LC-MS/MS analysis on the tissue samples, and plotted our results against standards and quadratic models in Supplementary Figures 2 and 3. Quadratic models are summarized in Supplementary Table 3.

**Figure 3.**
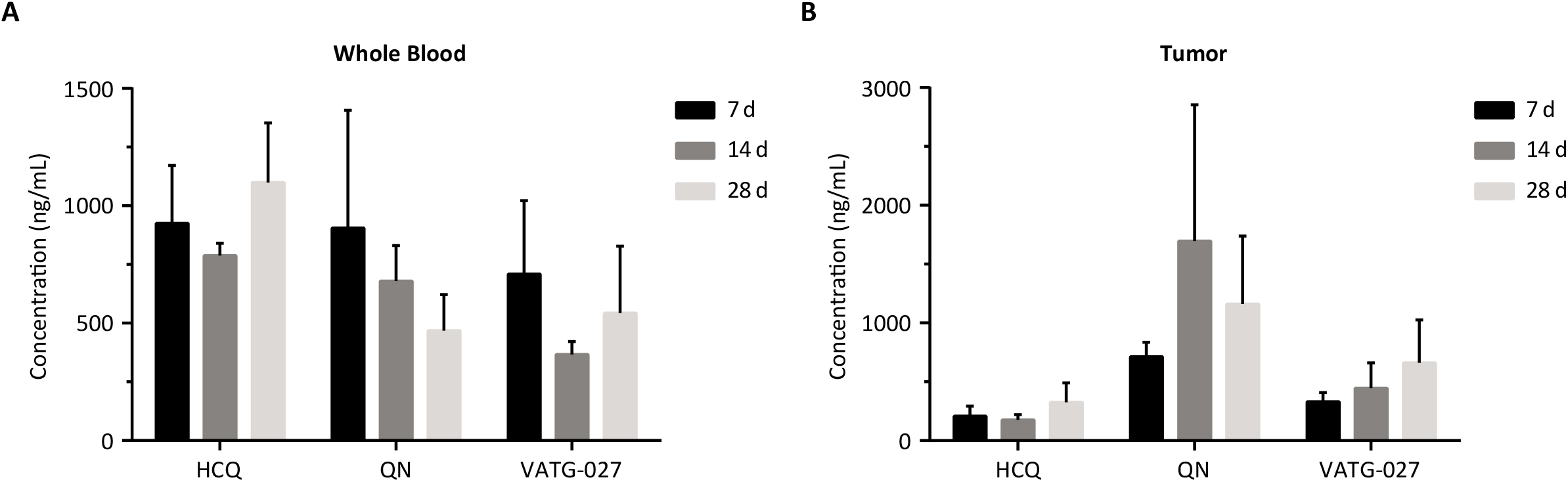
Whole blood and tumor concentrations over time. Athymic nude mice with subcutaneous A549 xenograft tumors were treated with 30 mg/kg HCQ, QN, or VATG-027 daily for 7, 14, or 28 d. 3 h after the last dose, (A) whole blood and (B) tumor samples were analyzed for compound concentrations by LC-MS/MS. QN and VATG-027 concentrations were compared to HCQ concentrations using one-way ANOVA; after 28 d of treatment, QN and VATG-027 whole blood concentrations were lower than HCQ whole blood concentrations (p<0.0001), and QN tumor concentrations were higher than HCQ (*p*=0.0002).

**Supplementary Figure 2. Standard curves for tumor tissues after 7 and 14 days of treatment.** Athymic nude mice with subcutaneous A549 xenograft tumors were treated with 30 mg/kg HCQ, QN, or VATG-027 daily for 7 or 14 d. Tissue samples were analyzed by LC-MS/MS for compound concentrations against an internal standard. Compound concentrations were calculated using quadratic models developed from the standard curves for each compound. Experimental samples/unknowns (red circles), standards (open circles), and the lines represent quadratic models generated from non-linear regressions.

**Supplementary Figure 3. Standard curves for whole blood and tumor tissues after 28 days of treatment.** Athymic nude mice with subcutaneous A549 xenograft tumors were treated with 30 mg/kg (A) HCQ, (B) QN, or (C) VATG-027 daily for 28 days. We analyzed tissue samples by LC-MS/MS to compare compound concentrations against an internal standard. For each compound, we used standard curves to generate quadratic models that determined compound concentrations. Tumor samples were analyzed by LC-MS/MS in two different batches, conducted on different days. Experimental samples/unknowns (red circles), standards (open circles), and the lines represent quadratic models generated from non-linear regressions.

Daily dosing of HCQ for 7, 14, or 28 d resulted in mean whole blood concentrations of 930, 790, and 1100 ng/mL, respectively (Figure 3A). Mean whole blood concentrations of QN were 910, 680, and 470 ng/mL after 7, 14, and 28 d of treatment, respectively (Figure 3A). Mean VATG-027 concentrations in whole blood were 710, 370, and 540 ng/mL after 7, 14, and 28 d of treatment, respectively (Figure 3A). After 28 d of treatment, whole blood concentrations of QN and VATG-027 were significantly different from HCQ concentrations (one-way ANOVA, *p*<0.0001). These data suggest that daily dosing of HCQ, QN, and VATG-027 does not cause accumulation in whole blood. Although generally comparable, these data also show that HCQ whole blood concentrations are higher than the acridine-based compounds QN and VATG-027 after 28 d of treatment.

Mean tumor concentrations of HCQ were 210, 180, and 330 ng/mL after 7, 14, and 28 d of treatment, respectively (Figure 3B). At these same timepoints, mean QN concentrations in tumor tissue were 710, 1700, and 1200 ng/mL, respectively (Figure 3B). VATG-027 mean intratumoral concentrations were 330, 450, and 660 ng/mL after 7, 14, and 28 d of treatment (Figure 3B). These data suggest that intratumoral concentrations of HCQ, QN, and VAG-027 may increase over time with daily dosing, although these differences were not statistically significant. In addition, these data show a general trend for acridine-based compounds achieving higher tumor concentrations than HCQ. Indeed, after 28 d of treatment, VATG-027 tumor concentrations were twice as high as HCQ concentrations, and QN concentrations were 4-fold above HCQ concentrations (one-way ANOVA, *p*=0.0002). When comparing whole blood and tumor concentrations, HCQ achieved both the highest whole blood concentrations and lowest tumor concentrations when compared to QN and VATG-027.

### Tissue distribution

To investigate the tissue distribution of HCQ, QN, and VATG-027, we treated athymic nude mice with 30 mg/kg of compound for 28 d. Because previous tissue samples were assayed 3 h post-gavage, we maintained the same experimental design for this study. We selected lung, liver, and kidney tissues as the primary clearance tissues of the body which often act as reservoirs of exogenous material. The tissues were compared to whole blood and tumor tissues as in previous studies. Tissue samples were analyzed by LC-MS/MS; experimental samples are plotted against standards and quadratic models in Supplementary Figures 3 and 4, and quadratic models are summarized in Supplementary Table 4.

As described in Figure 3, the mean whole blood concentrations of HCQ were 1100 ng/mL as compared to QN and VATG-027, which had mean whole blood concentrations of 470 and 540 ng/mL, respectively (Figure 4A, Table 2). QN and VATG-027 whole blood concentrations were significantly lower than HCQ whole blood concentrations (one-way ANOVA, *p*<0.0001). In addition, mean tumor concentrations after 28 d of treatment were 330, 1200, and 660 ng/mL for HCQ, QN, and VATG-027, respectively (Figure 4B, Table 2). Although both QN and VATG-027 achieved tumor concentrations higher than HCQ, only QN tumor concentrations reached statistical significance (one-way ANOVA, p=0.0002).

**Supplementary Figure 4.**
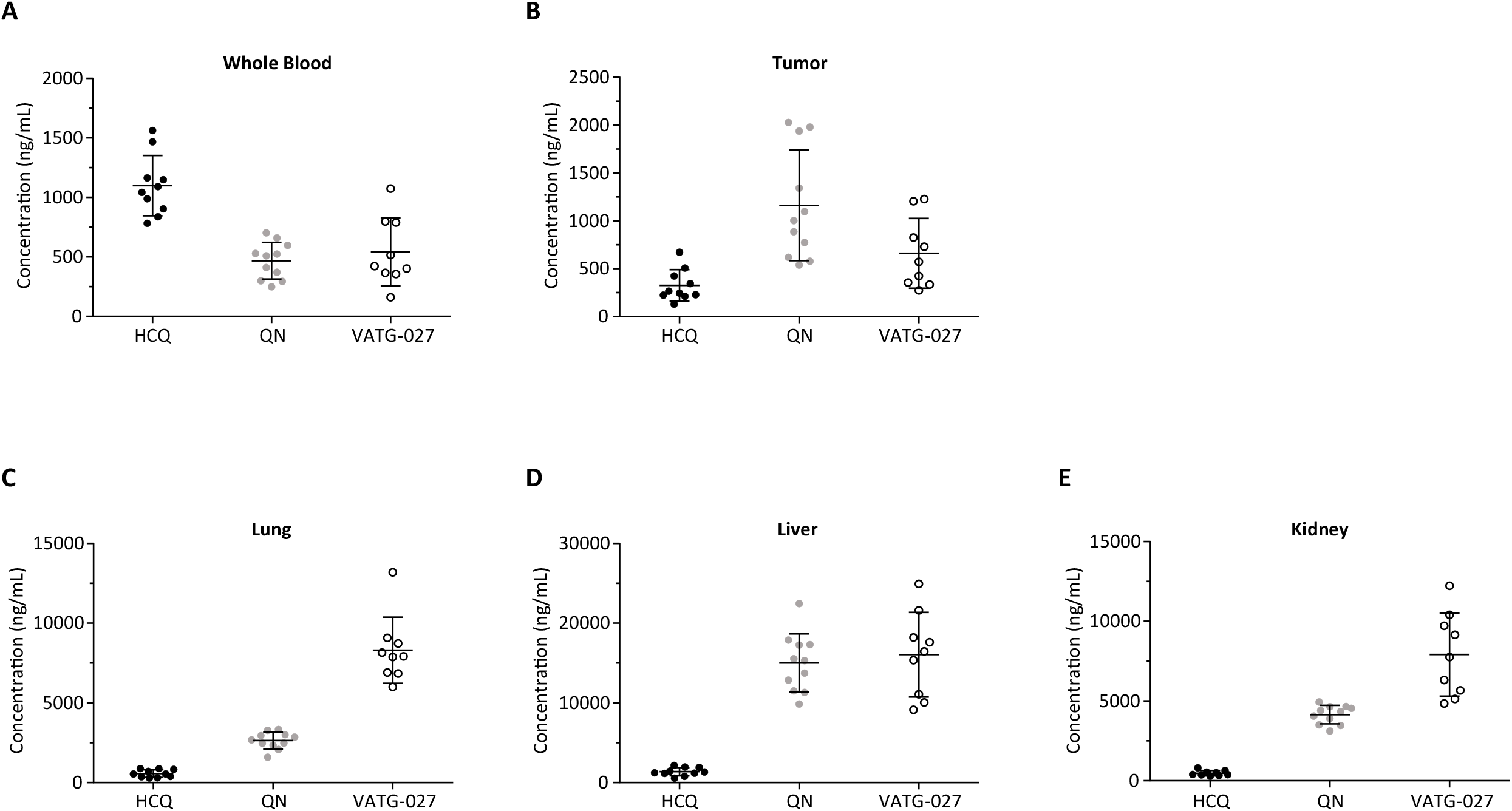
Standard curves for lung, liver, and kidney tissues after 28 days of treatment. Athymic nude mice with subcutaneous A549 xenograft tumors were treated with 30 mg/kg (A) HCQ, (B) QN, or (C) VATG-027 daily for 28 d. Lung, liver, and kidney samples were analyzed by LC-MS/MS for compound concentrations against an internal standard. We established compound concentrations using quadratic models based upon the standard curves for each compound. Experimental samples/unknowns are depicted in the red circles, standards are depicted in the open circles, and the lines represent quadratic models generated from non-linear regressions.

**Table 2.**
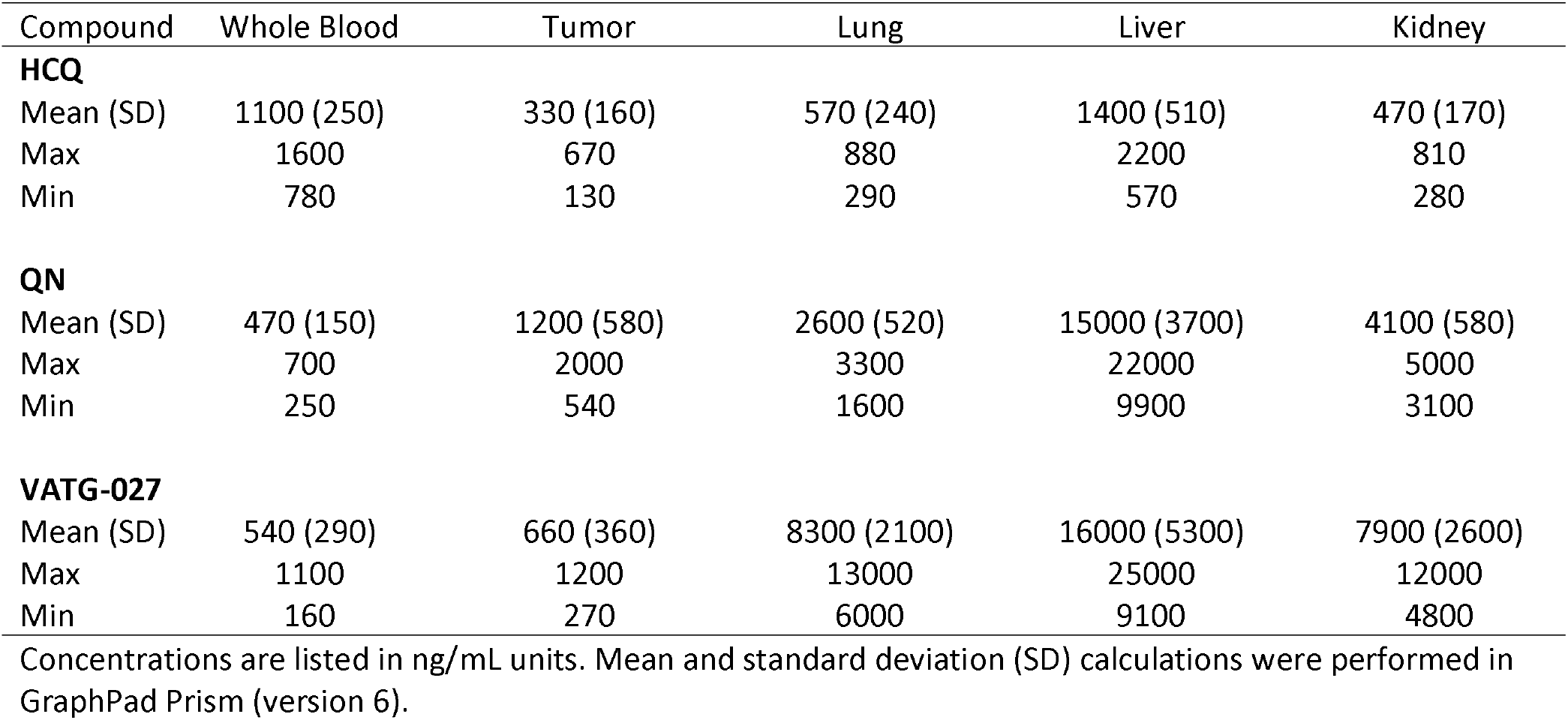
Tissue concentrations after 28 days of treatment.

After 28 d of treatment, the mean lung concentration for HCQ was 570 ng/mL (Figure 4C, Table 2). In contrast, QN and VATG-027 mean lung concentrations were 2600 and 8300 ng/mL, respectively (one-way ANOVA, QN vs. HCQ *p*=0.0009, VATG-027 vs. HCQ, *p*<0.0001). In the liver, mean HCQ concentrations were 1400 ng/mL, compared to 15000 and 16000 ng/mL for QN and VATG-027, respectively (Figure 4D, Table 2). Both QN and VATG-027 mean concentrations in the liver were significantly higher than HCQ concentrations (one-way ANOVA, *p*<0.0001). Mean kidney concentrations for HCQ, QN, and VATG-027 were 470, 4100, and 7900 ng/mL, respectively (Figure 4E, Table 2). Both QN and VATG-027 mean kidney concentrations were significantly higher than HCQ concentrations (one-way ANOVA, p<0.0001).

The tissue distribution data revealed HCQ mean concentrations increasing in the following order: subcutaneous tumor<kidney<lung<whole blood<liver (Table 2). QN mean concentrations increased as follows: whole blood<subcutaneous tumor<lung<kidney<liver (Table 2). VATG-027 tissue concentrations increased in the following order: whole blood<subcutaneous tumor<kidney<lung<liver (Table 2). These data indicate that the 4-aminoquinoline HCQ distributes into tissues uniquely when compared to the acridine-based compounds QN and VATG-027.

## DISCUSSION

In addition to being used as a treatment for malaria and autoimmune diseases, HCQ is being investigated as a lysosomotropic agent and autophagy inhibitor in cancer clinical trials (1, 6, 10). Clinical use of HCQ is limited given its adverse side effects and limited potency, as measured by autophagy inhibition in blood samples (30, 32). In response, we developed VATG-027, an acridine-based compound with improved potency over HCQ in multiple cell types (Supplementary Figure 1) (39, 40). In this first *in vivo* study, we report that VATG-027 is orally available at 30 mg/kg, and safe and tolerable when taken daily. Our observations support continued investigation of acridine-based compounds as lysosomotropic agents for the treatment of cancer, malaria, rheumatoid arthritis, and systemic lupus erythematosus.

In this report, we also provide a novel LC-MS/MS method for analyzing lysosomotropic compounds in various tissue types including whole blood, subcutaneous tumors, lungs, livers, and kidneys. Our method employs reversed-phase HPLC (Table 1) and tandem mass spectrometry to identify and quantitatively measure compound concentrations to a LLOQ of 2.5 ng/mL *(see METHODS).* Recently, Chhonker and colleagues published a LC-MS/MS method for the quantification of HCQ (43), again suggesting that LC-MS/MS methods are suited for quantifying 4-aminoquinoline and acridine compounds.

Chhonker *et al* and others have reported a high red blood cell to plasma partition coefficient for HCQ, suggesting that whole blood concentrations exceed plasma concentrations (43, 44). To accurately quantify the concentrations of HCQ, QN, and VATG-027 in the blood, we obtained whole blood samples rather than plasma samples. Although our data more accurately represent the compound concentrations in the blood, they may not reflect the concentrations of free compounds in the blood. Indeed, significant proportions of HCQ bind to hemoglobin and other proteins in whole blood (44, 45). These studies also support the lysosomotropic nature of HCQ causing uptake into lysosomes (4, 38). As such, use of whole blood is important to note when comparing results to other studies. Barnard and colleagues reported higher tumor concentrations of HCQ when compared to plasma (27). Barnard *et al.* use of plasma as opposed to whole blood in their study may explain this discrepancy.

The overall tissue distribution data suggests unique tissue concentrations between HCQ, QN, and VATG-027. Interestingly, HCQ concentrations in lungs, livers, and kidneys were significantly lower than QN and VATG-027 concentrations in the same tissues (Figure 4, Table 2). Of the tissues investigated, HCQ concentrations were highest in whole blood and liver. Similarly, QN and VATG-027 concentrations were highest in the liver. Elevated liver concentrations are expected from compounds delivered orally and experiencing the first pass effect (46). Interestingly, QN and VATG-027 concentrations were also significantly higher than HCQ in the lungs and kidneys, with VATG-027 concentrations being the highest of all three compounds (Figure 4, Table 2). These data suggest that acridine-based compounds preferentially accumulate in the lungs and kidneys when compared to the 4-aminoquinoline HCQ. As acridine scaffolds are used to develop next-generation lysosomotropic agents and autophagy inhibitors, it will be important to monitor tissue distribution and associated adverse side effects.

The acridine-based QN and VATG-027 also showed increased tumor concentrations over HCQ (Figure 3, Figure 4, Table 2). These data suggest that acridine-based autophagy inhibitors may have improved tumor uptake and efficacy over alternative chemical scaffolds. This may be especially true for lung and kidney cancers as QN and VATG-027 concentrations were highest in these tissues. Overall, our report provides a novel LC-MS/MS method and lysosomotropic compound (VATG-027) to further investigate pharmacokinetics and pharmacodynamics for the treatment of cancer, malaria, and auto immune disease.

## MATERIALS AND METHODS

### Animal Studies

All animal studies were approved by the IACUC at Van Andel Research Institute prior to beginning work; studies were conducted under IACUC protocols AUP-16-05-018 and XPA-17-01-004. For the single dose study, athymic nude, male mice were weighed the day before treatment and average weights were used to formulate autophagy inhibitors at 30 mg/kg concentrations in 60% v/v phosphate-buffered saline (PBS), 30% v/v polyethyleneglycol (PEG-400) (Sigma 202398), and 10% v/v ethanol vehicles. Hydroxychloroquine (HCQ) and quinacrine (QN) were dissolved directly into this vehicle, while VATG-027 was dissolved at 10x concentrations in 100% ethanol, and then diluted to 1x in PBS/PEG-400 immediately prior to treatment. Compounds were delivered in 100 μL volumes by oral gavage.

At 3, 6, 24, 48, and 168 h after treatment, animals were transferred to a necropsy suite and given Avertin (tert-amylalcohol from Sigma 24048-6 and 2, 2, 2-tribromoethanol from Sigma T48402) intraperitoneally using a 27 G needle (BD 309623). After two minutes, animals were tested for reflexes using the tail-pull and toe-pinch methods. Once animals were unresponsive, retro-orbital bleeds were performed to collect whole blood samples into ethylenediaminetetraacetic acid (EDTA)-coated, blood collection tubes (Greiner Bio-One 450480 and BD 367844). Blood samples were inverted several times to prevent clotting and then placed on dry ice. Cardiac perfusions were then performed using DPBS (Gibco 14040-133) and a Harvard Apparatus perfusion pump (PHD 2000 Infusion) at a flow rate of 5 mL/min through a winged catheter (Terumo SV-27EL). Catheters were placed into the apex of the heart and moved accordingly until organs turned pale. Five milliliters total of DPBS was used to perfuse each animal. Following perfusion, necropsy of kidneys, lungs, and livers was performed. All tissues were fixed in 4% paraformaldehyde (ChemCruz sc-281692) for 24 h, followed by storage in 70% v/v ethanol (diluted in distilled water).

For the daily dose study, A549 cells were grown to 95% confluence in 15 cm dishes (Corning 430599). Cells were collected and resuspended in RPMI-1640 (Gibco 11875-093) supplemented with 10% v/v FBS (Corning 35-010-CV) at concentrations of 3.6×10^6^ cells per 100 μL. Subcutaneous tumors were implanted (3.6×10^6^ cells) into the right flanks of athymic nude male mice using 27 G needles.

Tumor growth was measured using external calipers every 3 days; when tumors reached 150 mm^3^ +/− 20 mm^3^ (approximately 3-4 weeks after implantation), animals were randomly enrolled into treatment groups. Animals were treated daily by oral gavage with 30 mg/kg of vehicle (35% v/v ethanol in distilled water), HCQ, QN, or VATG-027, dissolved in 100 μL total volume. Animals were treated for 7 d (3 animals per group), 14 d (3 animals per group), or 28 d (10 animals per group for vehicle and HCQ, 11 animals per group for QN, and 9 animals per group for VATG-027). Three hours after the last treatment, animals were removed to a necropsy suite for collection of blood, tumor, lung, liver, and kidney tissues, as described above.

### Cell Culture

A549 cells were purchased from the American Type Culture Collection (ATCC CCL-185) and maintained in RPMI-1640 medium (Gibco 11875-093) supplemented with 10% v/v fetal bovine serum (FBS) (Corning 35-010-CV). Cells were grown in 6-well plates (Corning 3506) from seeding densities of 4.0xl0^4^ cells per well at 37°C, 5% CO_2_, and 93% humidity.

### Cell Lysis

At experimental endpoints, cells were removed from incubators to ice and media was aspirated from plates. Cells were washed once with 1x PBS at pH 7.4. Cell lysis buffer (10 mM monopotassium phosphate [Fisher Scientific P285], 1 mM EDTA [Acros Organics 40977], 50 mM glycerol phosphate disodium salt hydrate [Sigma G6501], 50 mM sodium fluoride [Fischer Scientific S299], 10% v/v NP-40 [Sigma 74385], 2% v/v Brij-35 [Pierce 20150], 2% w/v sodium deoxycholate [Fisher Scientific BP349], 200 mM EGTA [Acros Organics 428570500], 10 mM magnesium chloride [Acros Organics 41341], 1 mM sodium orthovanadate [Fisher Scientific S454], 2 mM dithiothreitol [Sigma 43816], and 1 % v/v protease cocktail inhibitor [Sigma P8340]) was added to cells at 80 μL per well of a 6-well plate, and cells were scraped into microfuge tubes. Cell lysis was allowed to take place over 10 min on ice. Cell debris was pelleted by centrifugation at 13,400 rpm and 4°C for 10 min. Protein supernatants were transferred to clean microfuge tubes and used to determine protein concentrations using Bradford assays on an Eppendorf Biophotometer (6131–02070). Briefly, Bradford reagent was made by diluting Bio-Rad (5000006) 1:5 in sterile, distilled water. Protein concentrations were determined by absorbance at 280 nm against a lysis buffer control, from 2 μL cell lysate samples diluted in 498 μL Bradford reagent in a plastic cuvette (Bio-Rad 223-9955). Instrument calibrations were performed monthly using standard curves of bovine serum albumin (Thermo Scientific 23209); calibrations were accepted with coefficient of variation percentages less than 5%.

### Cell Treatments

A549 cell treatments were performed diluted in fresh, full growth medium applied at the time of treatment. Hydroxychloroquine sulfate (Calbiochem 509272) and quinacrine dihydrochloride (Calbiochem 551850) were diluted in distilled water, while VATG-027 (Pharmaron PH-TGD-035-RS1) was diluted in DMSO. Inhibitor stocks were stored as single-use aliquots at -20°C. Vehicle treatments were performed at volumes equivalent to the highest volume of inhibitor applied to cells; DMSO vehicle treatments did not exceed 1% v/v.

### CellTiter-Glo

Cell viability assays were performed using CellTiter-Glo kits (Promega G7570). Cells were seeded at 1,000 cells per well into white-wall, clear-bottom 96-well dishes (Greiner Bio-One 655098). Twenty four hours after seeding, media were aspirated from wells and cells were treated with fresh media supplemented with vehicle, HCQ, QN, or VATG-027. Twenty four hours after treatment, media were aspirated from wells and cells were washed three times with 1x PBS. CellTiter-Glo reagent (prepared according to the manufacturer’s instructions, mixing the reagent and buffer solutions) was mixed 1:1 with 1x PBS and then applied to wells at 100 μL volumes. Plates were then shaken at room temperature and 550 rpm for 10 min to facilitate cell lysis, using a microplate shaker. After 10 min, luminescence was measured on a BioTek Synergy HT plate reader. Relative luminescence units (RLU) were normalized to the appropriate vehicle and data were constructed into figures using GraphPad Prism (version 6).

### Immunoblotting

Electrophoresis was performed in SDS-PAGE running buffer containing 25 mM Tris base, 192 mM glycine (Fisher Scientific BP381), and 0.1% w/v SDS. Electrophoresis was performed at 100 V for 2 h. After electrophoresis, proteins were transferred onto methanol-activated polyvinylidene fluoride (PVDF) membranes (Millipore IPVH00010) with 0.45 μm pores in transfer buffer at pH 8.3, containing 25 mM Tris base and 192 mM glycine. Transfers were performed at 4°C and either 550 mA for 3.5 h or 220 mA overnight (i.e., 16-20 h). Following transfers, membranes were blocked for 60 min in 5% w/v milk, diluted in l× TBS-T (0.2 M Tris base and 1.5 M NaCl [Fisher Scientific BP358] at pH 7.6, supplemented with 0.05% v/v Tween-20 [Affymetrix 20605]); blocking was performing rocking at room temperature.

After blocking, membranes were rinsed in l× TBS-T and cut according to the molecular weights of target proteins. Membranes were then incubated in primary antibodies (1:1,000 LC3B [Sigma L7543] and 1:10,000 β-actin [CST 3700]), diluted in 5% w/v bovine serum albumin (Sigma A7906) diluted in 1x TBS-T. Primary antibody incubations were performed rocking at 4°C overnight (i.e., 16-20 h). After primary antibody incubations, membranes were washed three times, rocking for 5 min each at room temperature, in 1xTBS-T.

Membranes were then incubated in ECL-compatible, horseradish peroxidase-conjugated secondary antibodies (anti-rabbit [GE Healthcare LNA934V], anti-mouse [GE Healthcare LNA931V]), diluted 1:5,000 in 1x TBS-T. Secondary antibody incubations were performed rocking at room temperature for 60 min. After secondary antibody incubations, membranes were washed three times, rocking for 5 min each at room temperature, in 1× TBS-T. Immediately before imaging, membranes were rinsed once in 1x PBS. Membranes were then arranged in a film cassette and transferred to a dark room for exposure to film (Denville Scientific, Inc. E3018). Films were processed using a Kodak X-OMAT 2000A film processor and electronically scanned.

### Mass Spectrometry

Tissues were removed from -80°C storage and placed on dry ice. In a petri dish (VWR 25384-326), a small portion of tissue was separated from the organ/tumor periphery using a razor blade and transferred to a microfuge tube on ice. Tissues were then shipped to PharmOptima on dry ice for mass spectrometry analysis. Briefly, tissue segments were transferred to 2 mL Precellys tubes and weighed. Two to three ceramic (zirconium oxide), 2.8 mm beads were added to each tube, along with a volume of diluent (50% v/v acetonitrile, 50% v/v water, and 0.2% v/v trifluoroacetic acid [TFA]) 10-20 times that of the weight of the tissue. Tissues were then homogenized using a Precellys 24 homogenizer (Bertin Instruments EQ03119-200-RD000.0) at 5,500 rpm and a maximum temperature threshold of 10°C; three thirty-second homogenization pulses were used with 20-s pauses between each pulse. Homogenized tissues were then stored at -80°C until extraction for mass spectrometry.

Liquid chromatography tandem mass spectrometry (LC-MS/MS) was utilized to determine the concentrations of compounds in tissue samples. Twelve-point standard curves were used for each compound and an internal standard (quinacrine mustard dihydrochloride) was used in every compound standard and experimental sample to account for instrumental variability. Briefly, 50 μL aliquots of blank matrix (i.e., whole blood or vehicle-treated tissue), standards, and experimental samples were aliquoted into 96-well plates, to which 20 μL fresh internal standard (diluted in 50% v/v methanol, 50% v/v water, and 0.2% v/v formic acid) was added. Acetonitrile at 200 μL volumes was added to each well, plates were mixed for 1 min in a multi-vortexer, and then plates were centrifuged at 4,000 rpm and 4°C for 10 min. Supernatants at 150 μL volumes were transferred to 96-well plates pre-filled with 150 μL of 100% v/v water and 0.2% v/v TFA.

Plates were then loaded onto an Agilent 1100 LC system. High performance liquid chromatography (HPLC) was performed using a Phenomenex Luna C18 (2) column. A gradient of two mobile phases was used for separation, over a total time of 5 min and using a flow rate of 0.5 mL/min (Table 1). Needle washes were performed using 100% v/v acetonitrile and 0.1% v/v formic acid. Immediately following separation by HPLC, samples were ionized and mass to charge ratios were measured using MS/MS on an Applied Biosystems/MDS Sciex API 4000 instrument.

Experimental sample peak areas were normalized to internal standard peak areas, generating a peak area ratio parameter used as the dependent variable for model fitting against a 12-point standard curve. Standard curves were run in duplicate and any calculated concentration that varied from the theoretical concentration by more than 20% was dropped from analysis. The lower limit of quantification (LLOQ) of the assay was 2.5 ng/mL. Quadratic models were used to fit the experimental data to the standard curves. Standard curve data are summarized in Figure 2 and Supplementary Figures 2, 3, and 4. Quadratic models are summarized in Supplementary Tables 2 and 3. Analysis was performed in GraphPad Prism (version 6).

### Statistics

Statistical analyses were performed in GraphPad Prism (version 6) using one-way ANOVA with an α value of 0.05. QN and VATG-027 mean values were always compared to HCQ mean values. Dunnet?s tests for multiple comparisons were used and *p* values represent adjusted *p* values for multiple comparisons.

### Tissue Lysis

Tissues were removed from -80°C storage and placed on dry ice. In a petri dish, a small portion of tissue was separated from the organ/tumor periphery using a razor blade and transferred to a microfuge tube on ice. Tissues were incubated in cell lysis buffer (see “Cell Lysis” section above) for 60 min on ice. Tissues were then sonicated using a Branson digital sonifier instrument and microtip at 30% amplitude for 5-s pulses until homogenous; tissues were not sonicated for more than four pulses total, and were not sonicated for more than two pulses sequentially. Homogenized tissues were centrifuged at 13,400 rpm and 4°C for 10 min. Protein supernatants were transferred to clean microfuge tubes and used to determine protein concentrations using Bradford assays, as described previously.

## Supporting information

## ACKNOWLEDGEMENTS

We thank members of the MacKeigan laboratory for critical discussions and feedback. We thank Yanli Su for assistance with animal husbandry and treatments. We thank David Bailey with PharmOptima, LLC for assistance with LC-MS/MS assay development. J.P.M. has research support from award number R01CA197398 from the National Cancer Institute, and a fellowship from the Van Andel Institute Graduate School to A.R.S. The funders had no role in study design, data collection and analysis, decision to publish, or preparation of the manuscript.

