## Supplementary figures and images for "Tissue distribution and tumor concentrations of hydroxychloroquine and quinacrine analogs in mice"

### Supplementary file 1

#### Supplementary Figure 1

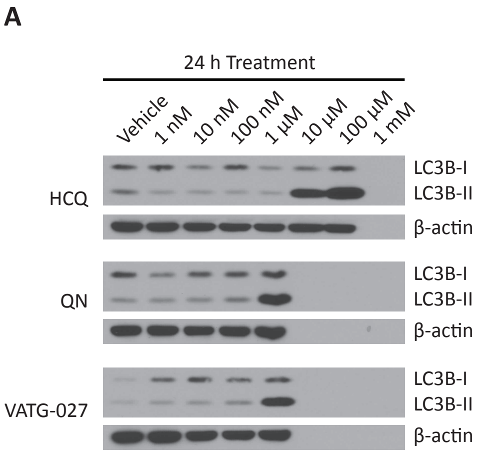


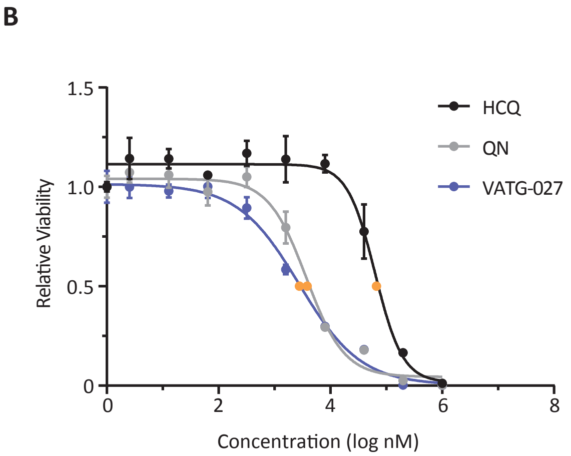

### Supplementary file 2

#### Supplementary Figure 2

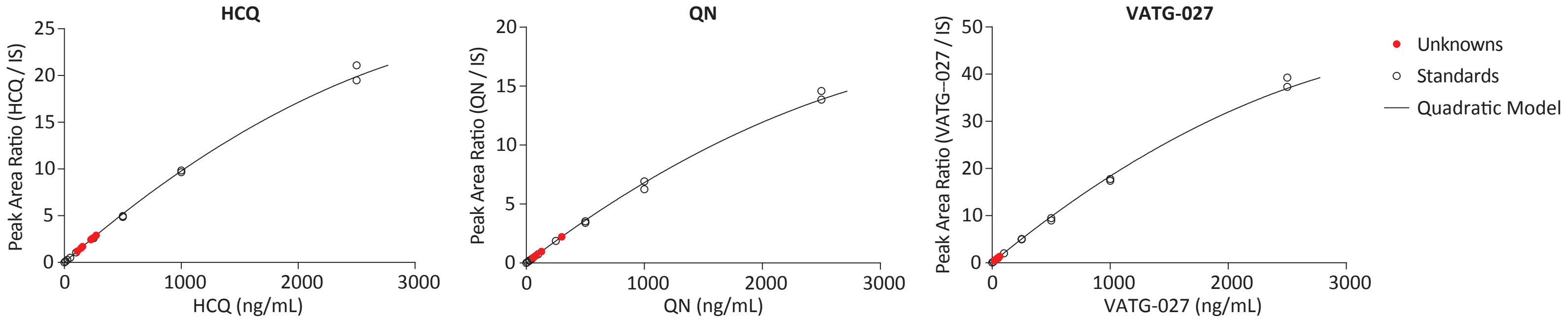

### Supplementary file 3

##### Supplementary Figure 3

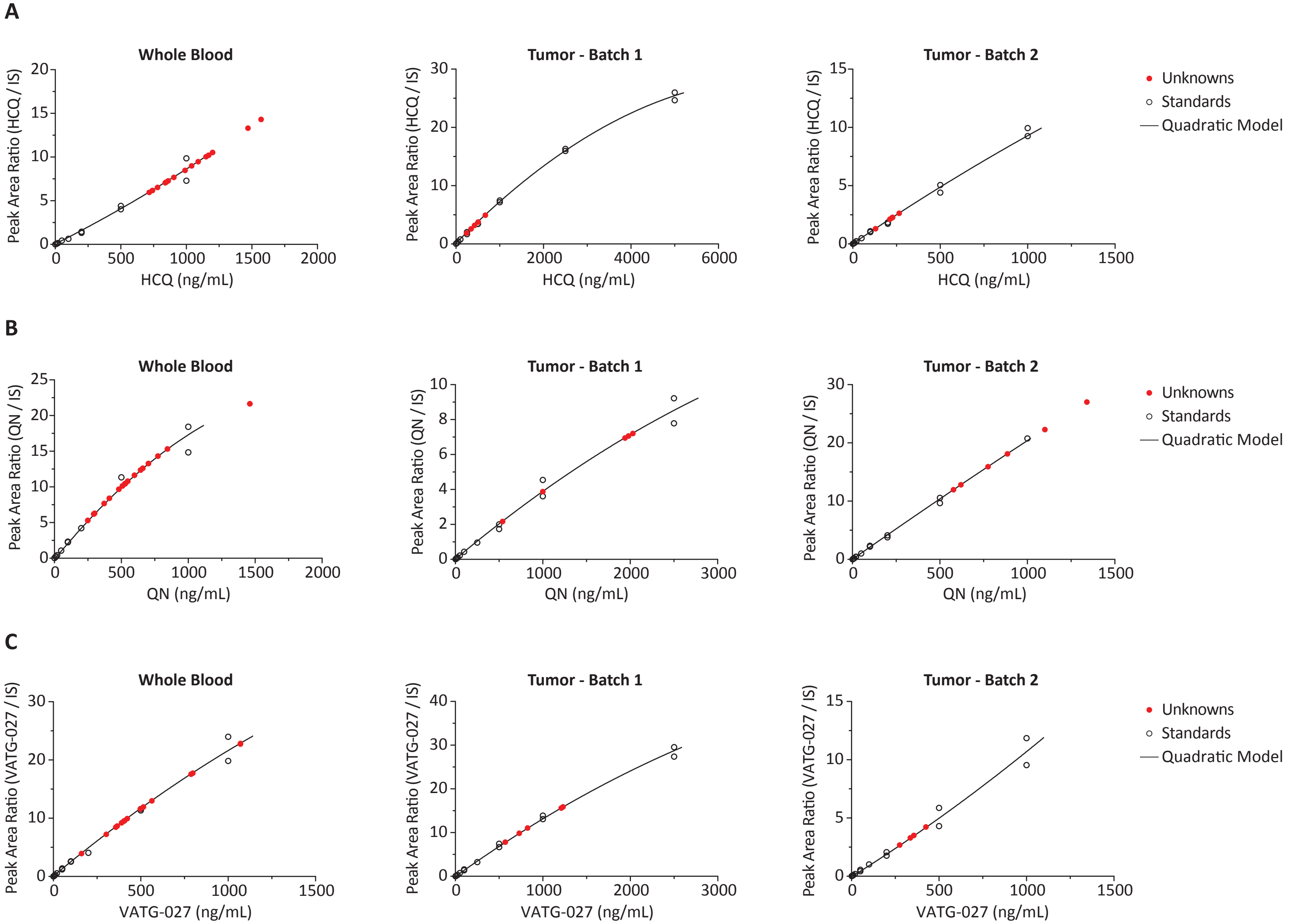

### Supplementary file 4

##### Supplementary Figure 4

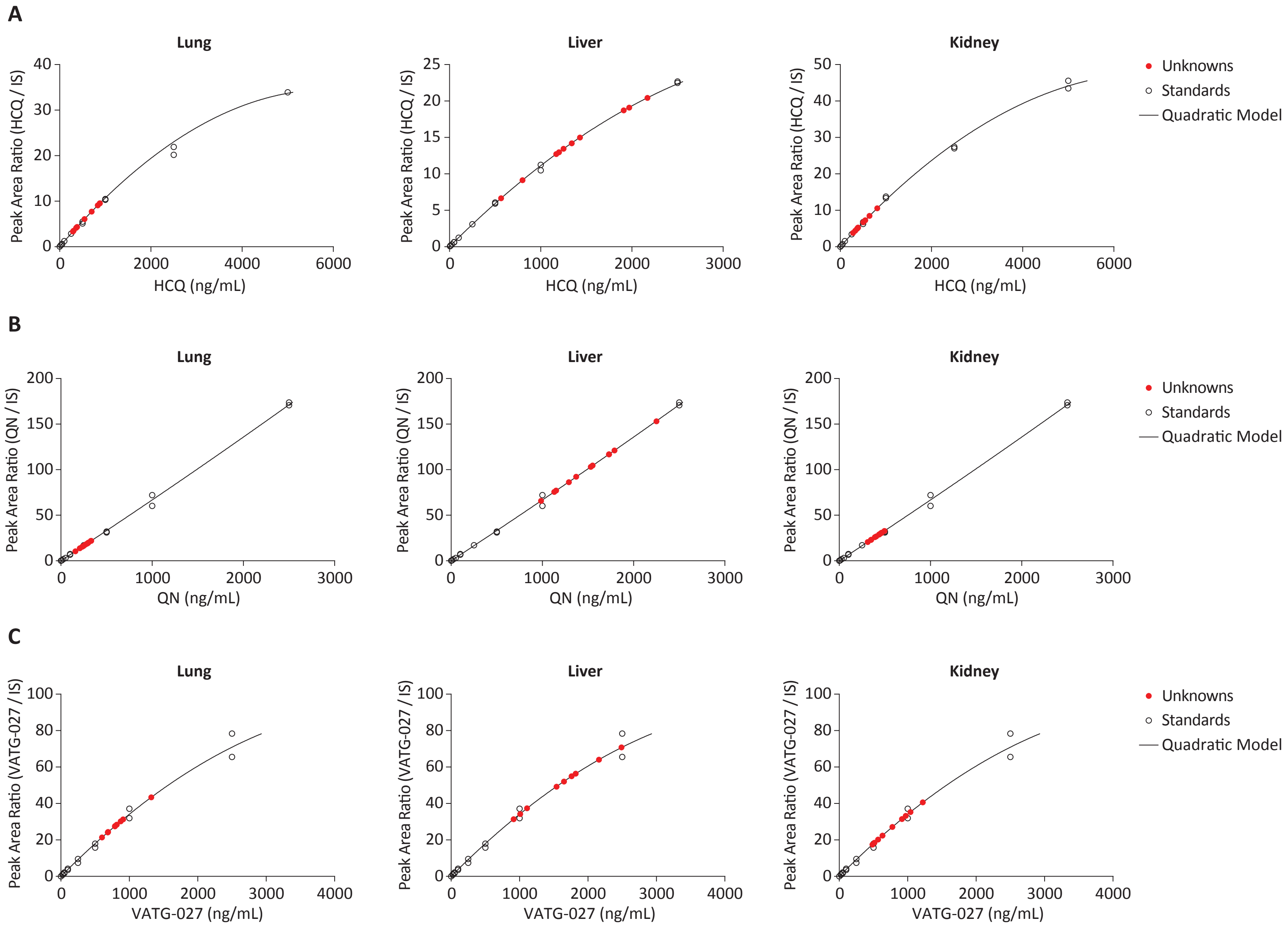
