## Supplementary material for "Tissue distribution and tumor concentrations of hydroxychloroquine and quinacrine analogs in mice"

**Supplementary Table 1. Physiochemical properties of compounds.**

| Property | HCQ | QN | VATG-027 |
|---|---|---|---|
| CAS Number | 118-42-3 | 83-89-6 | 1411646-03-1 |
| Molecular Weight (g/mol) | 335.87 | 399.96 | 412.96 |
| Water Solubility | ≤ 100 mM | ≤ 100 mM | Insoluble |
| DMSO Solubility | Insoluble | ≤ 10 mM | ≤ 100 mM |
| PPE Solubility | ≥ 41 mM | ≥ 38 mM | ≥ 43 mM |
| Ethanol Solubility | ≥ 20 mM | ≥ 19 mM | ≥ 21 mM |
| Color | White | Yellow | Yellow |
| Physical State | Solid salt | Solid powder | Solid powder |

Ethanol refers to 35% v/v 200-proof ethanol in water. Abbreviations include DMSO (dimethyl sulfoxide) and PPE (60% v/v phosphate-buffered saline, 30% v/v PEG-400, and 10% v/v 200-proof ethanol.
