## Supplementary material for "Tissue distribution and tumor concentrations of hydroxychloroquine and quinacrine analogs in mice"

**Supplementary Table 2. Quadratic models for single dose data.**

| Dose | Compound | Quadratic model |
|---|---|---|
| 30 mg/kg | HCQ | $y = -4.2*10^{-7}x^2 + 0.0041x - 0.00068$ |
| | QN | $y = 2.4*10^{-6}x^2 + 0.0015x + 0.0015$ |
| | VATG-027 | $y = -2.0*10^{-6}x^2 + 0.019x - 0.0010$ |
| 60 mg/kg | HCQ | $y = -4.2*10^{-7}x^2 + 0.0041x - 0.00068$ |
| | QN | $y = -1.0*10^{-6}x^2 + 0.019x - 0.00081$ |
| | VATG-027 | $y = -2.1*10^{-6}x^2 + 0.019x + 0.00050$ |
