## Supplementary material for "Tissue distribution and tumor concentrations of hydroxychloroquine and quinacrine analogs in mice"

**Supplementary Table 3. Quadratic models for daily dose data.**

| Tissue | Compound | Quadratic Model |
|---|---|---|
| Whole Blood | HCQ | $y = 1.0*10^{-6}x^2 + 0.0075x + 0.0015$ |
| | QN | $y = -5.3*10^{-6}x^2 + 0.023x + 0.0065$ |
| | VATG-027 | $y = -3.4*10^{-6}x^2 + 0.025x + 0.0017$ |
| Tumor (7 and 14 d) | HCQ | $y = -1.2*10^{-6}x^2 + 0.011x + 0.00022$ |
| | QN | $y = -8.4*10^{-7}x^2 + 0.0076x + 0.053$ |
| | VATG-027 | $y = -2.4*10^{-6}x^2 + 0.021x + 0.00024$ |
| Tumor (28 d, Batch 1) | HCQ | $y = -5.2*10^{-7}x^2 + 0.0077x + 0.016$ |
| | QN | $y = -3.1*10^{-7}x^2 + 0.0042x + 0.0023$ |
| | VATG-027 | $y = -1.1*10^{-6}x^2 + 0.014x + 0.0013$ |
| Tumor (28 d, Batch 2) | HCQ | $y = -8.9*10^{-7}x^2 + 0.010x + 0.0053$ |
| | QN | $y = -7.2*10^{-7}x^2 + 0.021x + 0.0063$ |
| | VATG-027 | $y = 1.4*10^{-6}x^2 + 0.0094x + 0.0033$ |
| Lung | HCQ | $y = -9.9*10^{-7}x^2 + 0.012x + 0.029$ |
| | QN | $y = 1.2*10^{-6}x^2 + 0.065x + 0.053$ |
| | VATG-027 | $y = -3.8*10^{-6}x^2 + 0.038x - 0.013$ |
| Liver | HCQ | $y = -1.5*10^{-6}x^2 + 0.013x + 0.017$ |
| | QN | $y = 1.2*10^{-6}x^2 + 0.065x + 0.053$ |
| | VATG-027 | $y = -3.8*10^{-6}x^2 + 0.038x - 0.013$ |
| Kidney | HCQ | $y = -10*10^{-7}x^2 + 0.014x + 0.0065$ |
| | QN | $y = 1.2*10^{-6}x^2 + 0.065x + 0.053$ |
| | VATG-027 | $y = -3.8*10^{-6}x^2 + 0.038x - 0.013$ |
